# What are the criteria for morphological cell death in *Dunaliella salina*?

**DOI:** 10.1101/2022.12.13.520199

**Authors:** Mahnaz Barmshuri, Bahman Kholdebarin, Saber Sadeghi, Zahra Faghih

**Author notes:** Correspondence, Co-Correspondence.

## Abstract

By finding morphological criteria for death in photosynthetic algal cells, one finds that the death of different populations of algae cells is manifested by various morphological changes. Present study, was undertaken to determine morphological criteria to be used in identifying cell death in unicellular green algae in their natural habitats. By applying the principles of formazan crystal formation due to MTT reduction in the presence of cells oxidoreductase enzymes, and the staining of saccharide complexes produced in photosynthesis by iodine reagent, morphological criteria were determined for cell death in *Dunaliella salina* collected from Maharloo lake and three different types of deaths were identified. Further studies have shown that these criteria can also be applied for fresh water algae and other taxon. Different ways of cell death in unicellular aquatic organisms can be used as monitoring tools for early warning of environmental hazards. We invite scientists, editors and reviewers to embark on establishing a much needed cell death classification committee for identifying different types of cell death and investigate mechanisms involved in unicellular aquatic algal cells.

**Significance Statement:** Staining with MTT and iodine reagents, are the best tools for distinguishing damages done to photosynthetic system in aquatic unicellular green algae following which cell death classification will be determined.

## Introduction

Green algae have essential roles in dynamic and structure of marine food network, biochemical carbon cycle and in bio-limiting elements in marine environments (1). It has been shown that, some eco-physiological processes of green algae present in marine ecosystems, affect both their death and the kind of death (2). Some evidences have been presented which indicate the presence of some factors affecting the controlled destruction of green algae and different forms of cells death which have been described as the algal responses to biological and non-biological stresses (3–7). Nevertheless, till now not only no systematic classification of different forms of deaths for unicellular green algae has been presented, but also due to the presence of common characteristics among photosynthesizing unicellular green algae, no uniform criteria have been defined for algal cell death. Unicellular green algae have common characteristics in reducing tetrazolium and also in staining of starch sheath by iodine around their rubisco granules. Using these common characteristics, we defined morphological criteria of cell death in *D. salina* algal. By determining the exact cause(s) of death, and by considering the criteria proposed by Nomenclature Committee on Cell Death (NCCD) the classification of cell death based on morphological criteria in unicellular *D. salina* and other algal taxa would be possible. Cell death in unicellular aquatic organisms occurs in different ways which can be used both as an early warning of environmental hazards and as a monitoring tool.

## Materials and Methods

### Cellular groups and staining of cells contents

The morphological changes and the possible biochemical pathways leading to cells death, were studied in four groups:

1. Cells collected from Maharloo Salt Lake.
2. Cells precipitated in cells suspension prepared from lake solution and kept for nine months under laboratory conditions without changing the suspension solution or taking subculture.
3. Cells treated with cis-platinum drug.
4. Cells exposed to visible light. The potential of tetrazolium salt reduction in photosynthetic electron transport system due to the presence of oxidoreductase enzymes in light reaction centers, and the staining of starch sheath around rubisco granules by iodine reagent, were used as the inner cellular criteria to study cells deaths. To determine the criteria for cells morphological death and its type, in each experiment more than 2 000 cells were used to be studied by Microscope Olympus (Japan) BX61 light microscope equipped with Microscope Digital Camara DP73 and 40 and 100 lenses.

### *D. salina* habitat and the site of its collection

*D. salina* algal cells were collected from a hypersaline lake (Maharloo Salt Lake) located at 23 kilometers east of Shiraz, Iran, between two geographical points of 29°32’29”N, 52°44’17”E and 29°19’21”N 52°54’32”E. The area of the lake is about 250 km^2^ (Fig. 1). The chemical and physical properties of the lake have been described by Zmamanpoor and Darmipouran (8) and its sediments have been reported by Kazemi *et al* (9).

**Fig. 1.**
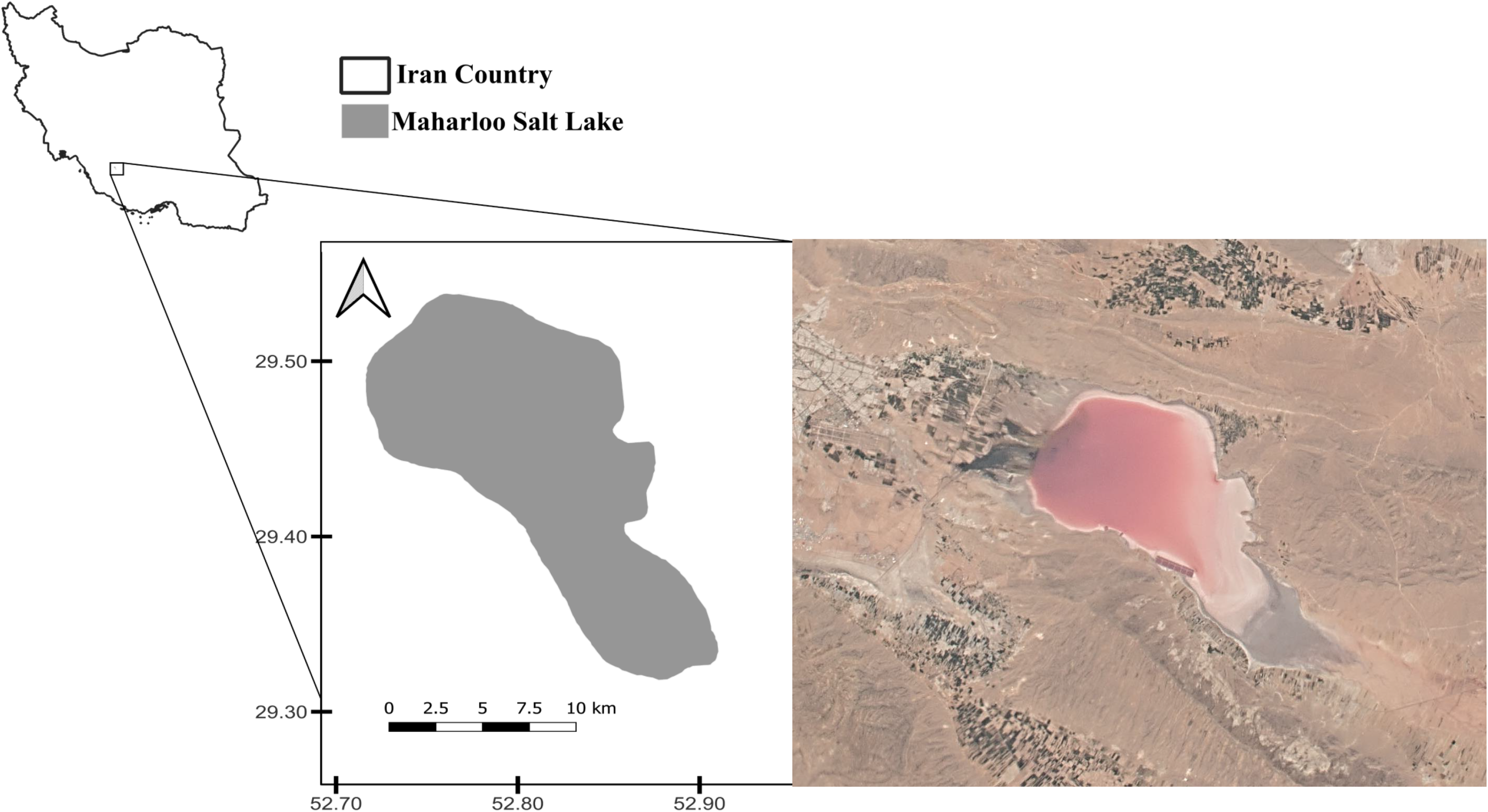
Geographical location of Maharloo Salt Lake shown in Iran map (Left). Aerial view of Maharloo Salt Lake. This picture has been taken by NASA International Space Station in 2015 (Right).

### Identifying *D. salina* and the conditions it was kept

The *D. salina* cell was identified using keys given by Pol *et al*, (10) and by Oren (11).

The cells suspensions collected from the lake, were transferred to the laboratory and kept at 25±1°C and at the irradiance of 40-50 μM photons m^-2^ s^-1^ supplied by cool white fluorescent lamps on a 16h:8h light/dark cycle (12).

### Inducing cell death by cis-platinum drug and by visible light

#### Death induced by cis-platinum drug

100 μL of *D. salina* cell suspensions, containing 1×10^5^ cells/mL, were added to a 96-well plate in triplicates and incubated at room temperature for 24 hours. Then, 100 μL cis-platinum (Mylan,1mg/ml) were added to each well and kept for 24 hours at 37±°C. Three wells containing 200 μL cells suspension were left untreated and were considered as negative controls.

### Cell death induction by visible light

When *D. salina* cells fluorescence was studied by visible light radiation produced by Olympus BX61 using barrier filter (GBA590), cells death was induced.

An experiment with 25 replications, was carried out in which algal cells were irradiated by visible light. A glass slide (24mm × 60mm) with 200 μL cell suspension containing 1×10^5^ cells/mL on it and covered with cover slip was prepared. Different sites of the slide were irradiated with visible light for two minutes produced by Olympus Bx61 microscope equipped with barrier filter (BA 590) and a dichromic mirror (DM 570). After irradiation, slides were washed with Ben-Amotz cell free culture solution (13). In order to have proper number of cells to study after irradiation, the cells in suspensions were sedimented by centrifuging at 3000 rpm for one minute. The supernatants were discarded and 2 mL Ben-Amotz culture solution were added to the sediments. After shaking slowly, the cells were transferred to 96-well plates for staining study. The moments in which algal cells and zygotes death occurred were recorded.

### Pyrenoids staining with iodine reagent

Pyrenoid staining was carried out by iodine reagent using 0.3gr Potassium Iodide (Sigma Aldrich, CAS-7681-11-0) and 0.15gr Iodine crystals (Merck, CAS-7553-56-2) dissolved in 50 mL distilled water. To 400 μL of cells suspension containing 1×10^5^ cells/mL, 2 μL iodine reagent were added slowly until the cell suspension color changed to dark violet. At this time, the cells were precipitated. The suspension solution was gently shaken and one drop was smeared on each slide.

### MTT assay for determining cells active metabolic sites

The metabolic activity of the cells in all groups were assessed by their potential to reduce tetrazolium salt (MTT (3-(4,5-dimethylthiazol-2-yl)-2,5-diphenyltetrazolium bromide)). Two hundred millilitres of cell solution from each group were transferred to 90 plates wells. Ten μlitres of 1mg/ml MTT solution (10 mg of MTT (Sigma Aldrich, CAS-298-93-1) in 10 mL of PBS. were added to each well. After overnight incubation at 37°C, the cells morphologies were studied by Olympus light microscope.

## Results

By staining both cells active metabolic sites and starch sheaths around rubisco granules, distinct morphological characteristics were observed which can be used to distinguish dead or dying cells from the living ones (Tables 1 and 2).

**Table 1.**
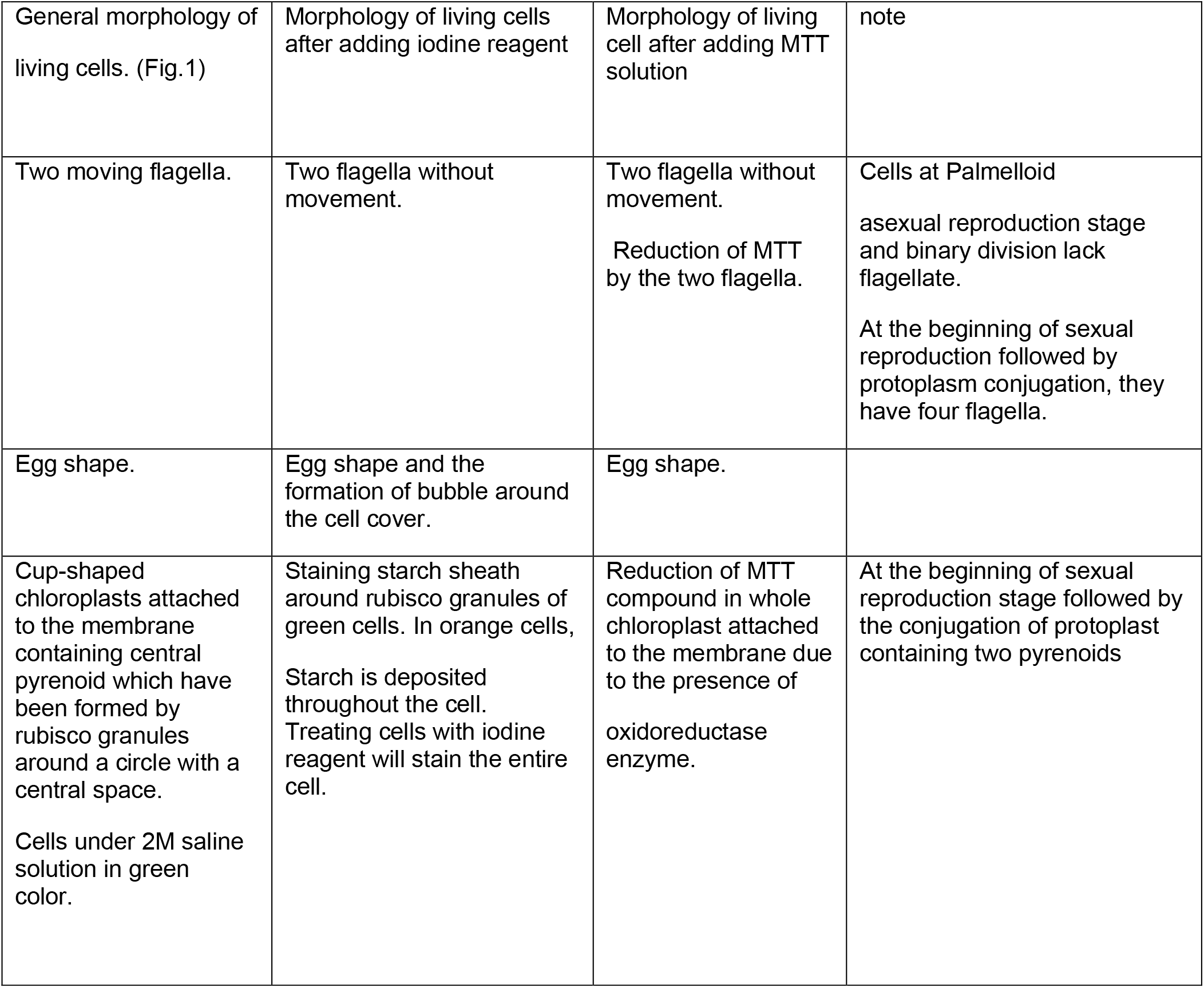
Morphological criteria to identify living cells in *D. salina* alga.

| General morphology of living cells. (Fig.1) | Morphology of living cells after adding iodine reagent | Morphology of living cell after adding MTT solution | note |
| --- | --- | --- | --- |
| Two moving flagella. | Two flagella without movement. | Two flagella without movement.<br>Reduction of MTT by the two flagella. | Cells at Palmelloid asexual reproduction stage and binary division lack flagellate.<br>At the beginning of sexual reproduction followed by protoplasm conjugation, they have four flagella. |
| Egg shape. | Egg shape and the formation of bubble around the cell cover. | Egg shape. |  |
| Cup-shaped chloroplasts attached to the membrane containing central pyrenoid which have been formed by rubisco granules around a circle with a central space.<br>Cells under 2M saline solution in green color. | Staining starch sheath around rubisco granules of green cells. In orange cells, Starch is deposited throughout the cell. Treating cells with iodine reagent will stain the entire cell. | Reduction of MTT compound in whole chloroplast attached to the membrane due to the presence of oxidoreductase enzyme. | At the beginning of sexual reproduction stage followed by the conjugation of protoplast containing two pyrenoids |

**Table 2.** Morphological characteristics of death cell in *D. salina*.

| General morphological characteristics of dead cell (Fig. 3). | Morphological specificity of starch sheath around rubisco granules (pyrenoid) stained by iodine reagent. | Morphological specificity of cells active metabolic sites stained by MTT solution* |
| --- | --- | --- |
| The morphology of dead cells found in Maharloo Salt Lake suspension solution (Fig. 3B3). |  |  |
| Egg shaped cells without flagella and mobility, disintegrated from the anterior side | Absence of starch sheath staining around rubisco granules. | Absence of staining of cells active metabolic sites. |
| Circular cells without flagellum and mobility. Depression along the cell length and complete separation of cell membrane from cell cover. | Presence of an irregular starch sheath around rubisco granules. | Absence of staining of cells active metabolic sites. |
| The morphology of dead cells induced by visible light (Fig. 3B1,B2) |  |  |
| Egg shaped cells without flagellum and mobility. Disrupted cell from anterior region | Absence of staining of starch sheath around rubisco granules. | Absence of staining of cells active metabolic sites. |
| The morphology of dead cell caused by cis-platinum drug. |  |  |
| Circular cell with two flagella without mobility. | Slight staining of starch sheath around rubisco granules. | Slight staining of active metabolic sites. |
| The morphology dead cells found in the sediment of old culture (Fig. 3 B3). |  |  |
| Compressed circular cells without flagella and mobility. Exudation of lipid droplets due to compression. | Staining of starch sheath around compressed rubisco granules. | Cells without staining of their active metabolic sites. Infection observed inside the cells. |
| Circular cells without flagella and mobility. Depression in part of the cell or complete separation of cell membrane from the cell cover. | Presence of starch sheath around rubisco granules in an irregular structure. | Absence of staining of cells active metabolic |

When is a cell alive? The morphological criteria used to identify *D. salina* living cells from the dead ones are presented in Table 1.

### When is a cell dead?

The Nomenclature Committee on Cell Death (NCCD), suggests that it is important to discriminate between dying as a process and death as an end point. To define cell death and related morphological characters, some unit criteria have been proposed (14). In the present study, the point-of-no-return has been considered as cell death, and according to NCCD suggestions, death in *D. salina* unicellular green alga has been classified. In green algae cells, if photosynthetic system as the most vital of cell system is damaged, photosynthesis will be stopped which in turn will be followed by cells death. When this happens, morphological changes will appear due to physiological changes taking place inside the cells. Based on the results obtained, cell deaths were classified in four groups. The morphological characteristics of dead cells in all studied groups are described in table 2

## Discussion

Studies of cell death in unicellular aquatic organisms have reflected different evidences (15–19).

Besides these evidences, we present morphological criteria to be used for defining death in studied algal cells. When serious damage is done to the reaction centers of photosynthetic organisms, the electron transfer systems and carbohydrate production will be stopped which will result in cell death. The electron transport chains operating in the light reaction phase of photosynthetic cells, are composed of oxidoreductase enzymes (20–25) that also cause the reduction of tetrazolium salt which is a staining method used to study the survival, reproduction and the metabolic activity in mammalian cells. MTT assay is used to study mitochondrial activity (26–28). In different studies done in aquatic unicellular organisms, MTT assay is used to investigate cells viability in response to different factors (29–33). The pyrenoid matrix which has been developed in photosynthetic organisms, is mainly a condensed package of rubisco (ribulose-1,5-bisphosphate carboxylase/oxygenase) enzyme which is the key enzyme in carbon dioxide reduction cycle of photosynthesis (Calvin cycle). The starch produced in photosynthesis, is deposited around rubisco granules (33–36). Iodine causes amylose present in starch to turn into dark blue color. In the presence of potassium iodide (IK), the iodine dissolved in water will make a complex soluble linear triiodide which then enters the starch coil and causes a very blue-dark color to appear (37).

The cell death caused by the application of anticancer drug cis-platinum which is attached to DNA (38), will block cell division and causes apoptosis (38–40). Directly or indirectly, it also increases ROS production in mitochondria (41–43) which is followed by the inhibition of respiratory system in mitochondria (44). Besides attaching to nuclear DNA, other causes of cis-platinum toxicities in the cells, are attaching to mitochondrial DNA, RNA, proteins and small peptides (45–49). The presence of semi-autonomous chloroplast DNA has been confirmed biochemically and by using electron microscope (50–53). The effects of cis-platinum on chloroplasts genome probably causes the inhibition of gene expression of oxidoreductase enzymes operating between the two light reaction centers. In green unicellular algae cells, MTT reduction is not solely dependent on mitochondrial activity. The absence of ATP and NADPH production, will cause a drastic reduction in algal cells potentials for MTT reduction and also in providing enough chemical energy for carbon dioxide reduction to produce carbohydrates and other organic substances by light independent phase of photosynthetic reactions.

The photosynthetic systems of green plants and green algae have two light reaction centers called photosystem I (P700) and photosystem II (P680) (54–56). A question which will arise is that, in spite of the proper interaction of these two light reaction centers by visible light spectra, why *D. salina* cells, when treated with visible light produced by microscope under laboratory conditions, undergo the process of cells death is induced? Also, why the morphologies of these types of dead cells are also observed in cells living in Maharloo Salt Lake? It has been clearly found out that the light dependent chemical reactions of photosynthesis are due to the presence of photosensitizers in the light reaction centers of photosynthetic organisms (57).

Upon absorption of light energy, chlorophyll in harvesting antenna complex attains a high energy but short-lived singlet excited state (^1^Chl*)(58). In photochemical quenching, part of this light energy absorbed in singlet excited state (^1^Chl^*^), is transferred to photosynthetic light reaction center (P680) to operate the photosynthetic electron transport chain (58). In the present study, by taking film from the moment death was induced by visible light, it was noticed that the cells chlorophyll molecules which were exposed to visible light, were not able to quench the unused light energy ^1^Chl^*^ by fluorescence (Film No.1). The unquenched light energy caused the production of triplet excited chlorophyll (^3^Chl^*^). In spite of the presence of high quantity of beta carotene molecules inside the cells, they were not able to quench ^3^Chl^*^. Probably, the high amount of ^3^Chl^*^ reacts with molecular oxygen (O2) produced by water-splitting reaction in the oxygen-evolving complex and will produce very unstable and reactive singlet oxygen (^1^O2) which causes the cells to be under oxidative stress (58–61).

It has been shown that the agents causing membrane disruption due to lipid oxidation in cancer treatment by photodynamic therapy, are the accumulation of both reactive oxygen species (ROS) and singlet oxygen species (SOS) which cause cells necrosis. Also, in the process of photodynamic therapy (PDT) which is based on the presence of a photosensitive compound that absorbs NIR light followed by a ligand-release reaction which results in micro-perforations of cell membrane that quickly coalesce into blebs followed by cells ruptures. Probably, by the release of photosynthesized ligands in chloroplasts, physical stresses and cell membrane disruption will be followed causing transmembrane osmotic gradient, the results of which is the cell to be contracted from anterior side toward cell center causing sudden cell rupture at the front side (Film No.2 and 3). Apparently, all evidences have indicated that in relation to cell death caused by NIR, can also be applied to cell death in salina alga cells by visible light. When visible light radiated by microscope barrier filter (GBA590) falls on glass slide, probably after passing through slide cover and reaching the slide surface, light interference phenomenon (62) will take place which increases the light intensity radiated by microscope causing cell membrane oxidation and destruction. The possible explanation for this type of algal cells death in the Moharloo Salt lake could be as follows. In the right conditions, the infrared spectrum of the visible light passing through the salt crystals (63) upon interference, achieving higher intensity, kills the trapped cells causing their necrosis type death. Results shown in our study and also in photodynamic therapy indicate that another way for recognizing death in cells having light reaction centers, is bleaching and the loss of fluorescence (64–69), (Fig. 3B–2). In algal cells containing photosensitizer (s) in their light reaction centers, cell bleaching caused by visible light, the light which is radiated after death is due to the phosphorescence of light reaction centers (Film No. 1). Different studies have confirmed the presence of pericellular matrix of glycoprotein type in *D. salina* alga (10, 70–73). The loss of cell protective cover affected by different factors, will cause death in these algal cells by different ways. Glycocalyx acts as an intermediate and protects cell membrane against direct physical forces and environmental stresses allowing it to maintain its integrity. The attachment of cell membrane to cell cover takes place by membrane proteins (74–75). *D. salina* alga which lives in hypersaline environments, adjusts itself to these ecosystems by cell plasma membrane protein kinases and also by internal glycerol synthesis which help cells to make osmotic adjustment to high external saline conditions (76). Damages done to membrane proteins will be followed by the disruption in glycocalyx action, the most important results of which is the disturbance of osmotic balance inside the algal cells. The loss of attachment between membrane and cell cover, probably disturbs the expression of membrane protein kinases causing cell dehydration and the separation of cell membrane and cell cover which results in cell death by plasmolysis in *D. salina* cells (Fig. 3A).

**Fig. 2.**
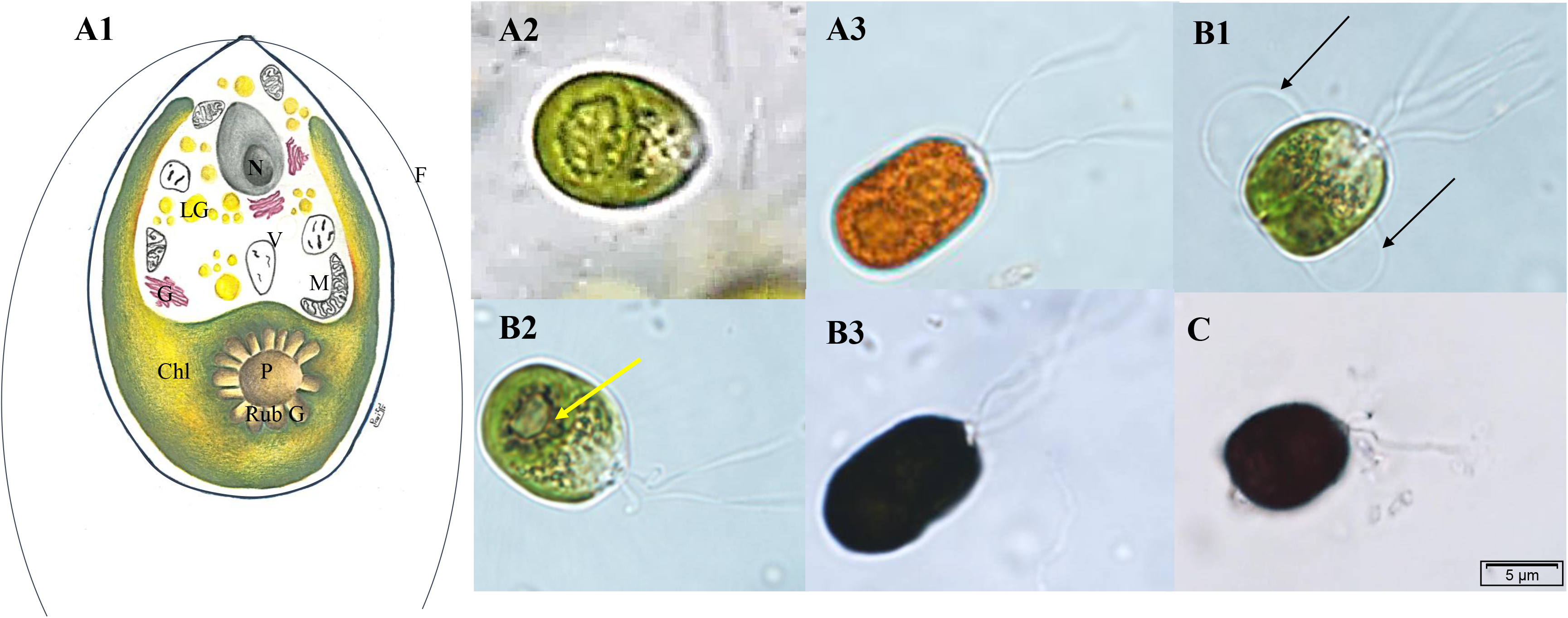
General morphology of living cells. **A1** Schematic diagram of a unicellular green *D. salina* cell. **F** flagella. **N** nucleus. **LG** lipid globules. **V** vacuole. **M** mitochondria. **Chl** chloroplast. **P** pyrenoid **Rub G** rubisco granules. **A2** Green alga cell. **A3** Cell under salinity stress which has produced beta carotene. **B1** *D. salina* alga egg cell (Hologamy) at the early stage of protoplast conjugation. Four flagella, two pyrenoids, production of membrane bubble after treating cells with iodine reagent (black arrows). **B2** Green algal cell stained with iodine reagent. Staining of starch plates around rubisco granules (white arrow). **B3** Staining of whole cell containing beta carotene. There is a direct relation between beta carotene and starch contents. We recommend this type of staining to determine the best time for beta carotene extraction from *D. Salina* algal cells in commercial cultures. **C** Algal cell stained with MTT. MTT reduction in the whole cup-shaped chloroplast of living cell.

**Fig. 3.**
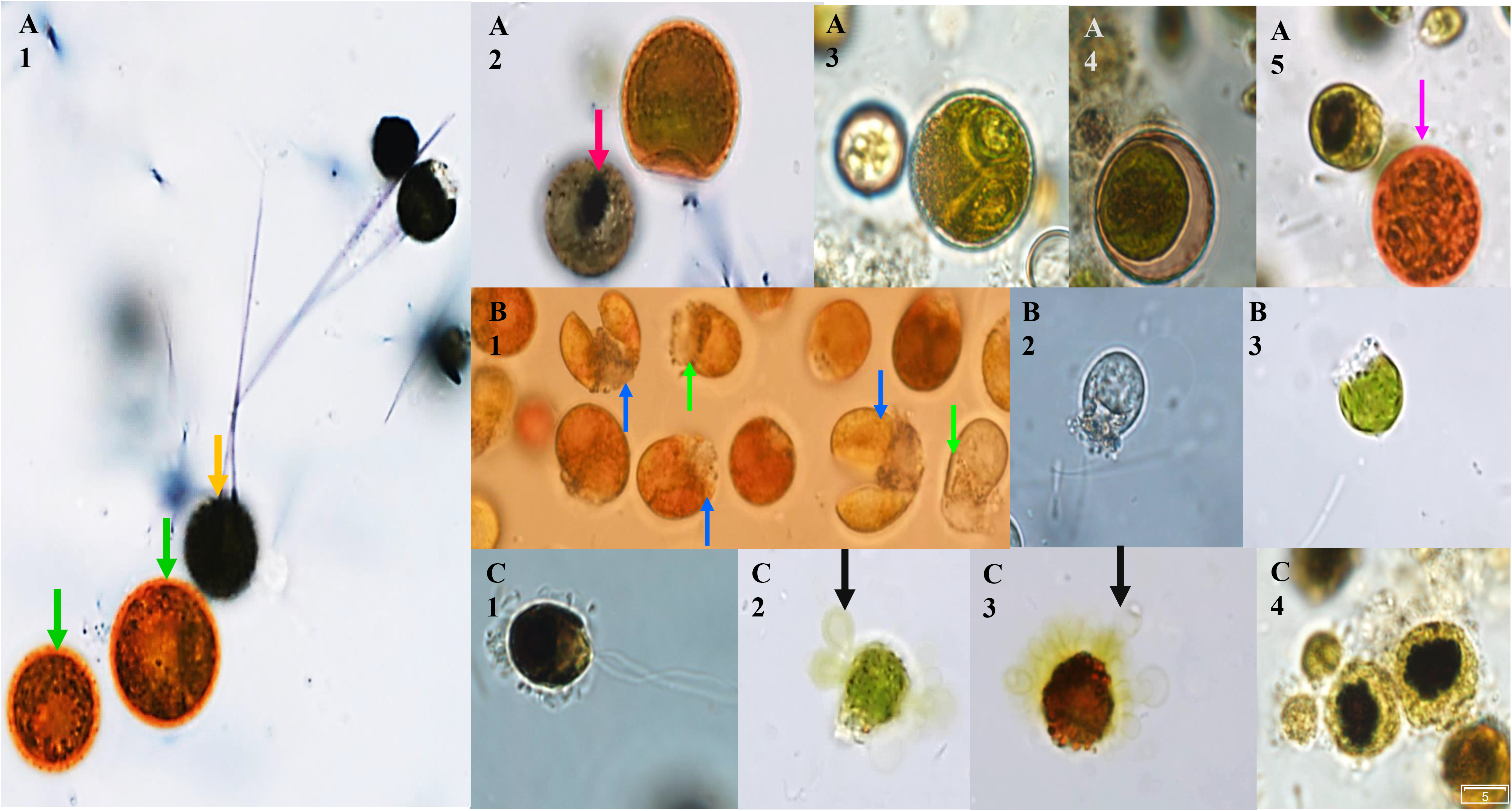
A. Morphology of *D. salina* cell death caused by plasmolysis. **A1** Incipient plasmolysis (at the early stage of Autosapore) in single algal orange cell which cannot reduce MTT (green arrow). Reduction of MTT in living cell chloroplast (blue arrow). Slight staining of cells active metabolic sites and detection of infection inside the cells (red arrows). **A2** Incipient plasmolysis in egg cell. **A3** Depression in cell due to cell volume reduction. **A4** Rupture of the connection between cell membrane and cell cover. **A5** The response of orange dead cell compared to living green cell to iodine reagent (purple arrow). **B. The morphology of *D. salina* cell died by necrosis. B1** Single cell death (at the early stage of Autosapore) by necrosis induced by visible light (green arrow) and algal egg cell (blue arrows). **B2** Cell color fading due to continuous radiation. **B3** Cell died by necrosis found in Maharloo lake. **C. Morphology of *D. salina* cell died by secretion. C1** Microbes have surrounded algal cell**. C2** and **C3** Disruption of osmotic equilibrium, depression of green and orange cells and lipid drops exudation from cells due to cells microbial infection. **C4** Pyrenoid compression.

In present study, another type of cell death was observed only in cells sedimented in old culture solutions. We believe this type of cell death is due to the loss of glycocalyx action caused by microbial infection of the algal cells. The high population of microbes and their secretions, cause the viscosity of suspension at the bottom of container to be increased. Low algal mobility at the bottom of culture container, will result more algal cells to be attacked by pathogens followed by the damage done to glycocalyx. As this happens, cells will not be able to control their osmotic potentials which causes the cells to lose their water content and they collapse all of a sudden. Because of the pores produced in both cell cover and cell membrane by microbial attacks, the lipids macromolecules will exude out of the cell. We use the term “secretion” for this type of cells death (Fig. 3C).

Since for the extraction of macromolecules from inside the cells, they should be lysed (76–77), and also for controlling algal bloom, this type of cell death can be used as a tool for both macromolecule extraction and controlling algal bloom (78).

In this study, by using two cell staining technics that is, with MTT and with iodine reagents, we present distinct morphological criteria to be used to define death in aquatic algal cells. Chloroplasts as the sites of photosynthetic electron transport chains, oxidoreductase enzymes and pyrenoids, are considered as the places for storing starch sheets produced as the product of photosynthetic carbon reduction cycle and were considered as the most vital sites to be used for these two vital staining technics. Although it may seem that these two staining technics are simple, they can be used for obtaining important and exact information about the function of photosynthesis in unicellular aquatic algal cells. This information, provided us with distinct criteria to define death for algal cells which was followed by showing us three types of non-physiological cell deaths.

## Acknowledgments

We would like to thank the Central Laboratory of Shiraz Medical School, Shiraz University of Medical Sciences, affiliated to Iran High-Tech Laboratory Network, for providing the Olympus microscope. We also thank the entomology laboratory of Shiraz University for providing travel and sampling expenses.

## Author Contributions

MB conceived the original idea, designed the methodology, supervised the project, performed the experiments and wrote the original draft, prepared and submitted the manuscript. BKH supervised the study, edited and revised the manuscript. SS supervised the study and provided travel and sampling expenses and with ZF edited the manuscript. All authors reviewed and approved the final manuscript.

## Conflict of interest

The authors declare that they have no competing interests in relation to this work.

## Funding

This work was supported by Fars Elites Charity Association.

